# Dysregulated RNA G-quadruplex binding proteins reveal shifted stress responses in Alzheimer’s Disease

**DOI:** 10.64898/2026.01.20.700564

**Authors:** Xingyu Zhou, Yuanyuan Zhang, Lei Sun, Chun Kit Kwok, Jilin Zhang

## Abstract

Single-cell sequencing has reshaped the research paradigm of Alzheimer’s Disease (AD) by revealing the heterogeneous transcriptional states of brain cells. Large single-cell cohort datasets, such as ROSMAP and SEA-AD, provide valuable atlases but primarily focus on RNA expression and chromatin accessibility, often overlooking the roles of RNA secondary structures. Although RNA G-quadruplexes (rG4s) are increasingly recognized as regulators of neurodegeneration, their interacting RNA-binding proteins (RBPs) remain poorly understood. We analyzed multiple independent scRNA-seq datasets to investigate rG4-associated biological functions across AD-relevant cell types and transcriptional states. We discovered that rG4-binding RBPs (RG4BPs) directly regulate essential glial cell functions, which are frequently impaired in AD. At the transcriptional state level, distinct sets of dysregulated RG4BPs correspond to the impaired state-specific biological features of astrocytes and microglia during AD progression, including a shift from an acute protective state to a chronic stress-associated state. This progressive state transition is accompanied by glial exhaustion and accumulated rG4s, which ultimately compromise glial support functions. We report several RG4BPs that are known regulators, including CIRBP, HSP90AA1, VIM, and PICALM, whose altered expression is associated with either pathological activation or severe functional impairment. These dysregulated RG4BPs provide a lens to examine the mechanism of rG4 accumulation in AD and position RG4BPs as novel targets for understanding and potentially intervening in AD progression.

## Introduction

Alzheimer’s disease (AD) is the major cause of dementia, which became the 7^th^ leading cause of death in 2021^1^. Given its global health impact, drugs targeting amyloid-beta (Aβ) plaques, like lecanemab ^2^ and donanemab ^3^, have been approved. However, their limited efficacy and adverse effects underscore a need to better understand underlying molecular drivers of AD ^4^. Although none of the existing hypotheses comprehensively identify the molecular drivers of sporadic AD onset, RNA metabolism has emerged as a critical axis in neurodegenerative disorders, implicating RNA-binding proteins (RBPs) and their target RNAs as potential pathogenic factors. Multiple lines of evidence indeed characterize dysregulated RBPs as detrimental regulators ^5–7^, but their detailed mechanistic roles in regulating RNAs remain poorly understood.

Technical advances in single-cell sequencing have revolutionized the scale and resolution of examining AD pathogenesis, enabling an in-depth dissection of dysregulated genes in transcriptionally heterogeneous cell populations sharing the same cell identity. Several studies have demonstrated that gene expression dynamics across cell populations are critical to maintain physiological haemostasis, and their dysregulation is directly associated with pathological conditions ^8–13^. Indeed, the expression of many genes across these transcriptional states becomes dysregulated in AD astrocytes and microglia, implying their indispensable regulatory roles during AD onset and progression ^13,14^. In astrocytes, the transition from a homeostatic to a reactive state could be hijacked by shifting into a metabolic state rather than progressing to an intermediate state. These rewired transcriptional state transitions lead to astrocyte exhaustion due to failed proteolysis and impaired energy metabolism ^15^. In microglia, the altered expression profiles of disease associated microglia further highlight the systematically impaired cellular functions ^13^. In response to the physiological changes in AD brains, such as increased phosphorylated tau levels and Aβ plaque accumulation, dysregulated state transitions can profoundly alter the biological functions of glial cell populations, thereby reshaping the neuronal microenvironment. Molecular characteristics from these unique profiling studies highlight the central role of transcriptional state in cells sharing the same identity during AD progression. While an elevated oxidative stress response is an AD hallmark ^16^, oxidative stress can also alter RNA folding substantially, suggesting that RNA secondary structures may participate in stress-associated transcriptional state dysregulation^17^. Nonetheless, structural RNA elements, such as RNA G-quadruplexes (rG4s), have not been systematically investigated in this context. Thus, the roles of RNA secondary structures and their underlying mechanisms during transcriptional state progression remain elusive.

The non-canonical RNA secondary structure rG4 has been shown to exacerbate cellular senescence by mediating ribosome pausing in the mouse brain ^18^, and to coordinate stress-granule assembly that regulates synaptic function in neurons of the mouse forebrain ^19^. Moreover, rG4 is likely accumulated in both neurons and astrocytes (both RNA and DNA G4), indicating that rG4 represents an indispensable RNA structural layer in the central nervous system ^20,21^. Recent studies indicate that rG4s and rG4 binding proteins (RG4BPs), including alpha-synuclein (SNCA), exhibit a positive correlation with AD severity and progression ^17,22^. In the meantime, lithium ions are trapped in Aβ plaques, leading to an imbalanced cation homeostasis in AD ^23^. Because lithium destabilizes rG4 structures, its sequestration in Aβ plaques may directly influence RNA structural dynamics and ion homeostasis under chronic stress conditions. While lithium supplement alleviates functional decline in AD models, the underlying molecular mechanisms and relevance to AD etiology remain unclear. Together, these observations suggest that dysregulated rG4 dynamics may link ion imbalance, stress responses, and transcriptional dysfunction in AD. Therefore, charting rG4 dynamics and RG4BPs can facilitate the functional dissection of AD pathogenesis. In this study, we combined experimentally confirmed

RG4BPs with public single-nucleus RNA sequencing (snRNA-seq) datasets to investigate the rG4-interacting RBPs, with a particular focus on their cell-type- and state-specific dysregulation. Complemented by independent scRNA-seq datasets ^24^, proteomic evidence ^25^, and RBP binding profiles ^26^, our findings establish dysregulated cellular stress responses mediated by rG4–RG4BP interactions as a plausible molecular mechanism linking multiple AD hallmarks.

## Results

### Altered expression of rG4-interacting RNA-binding proteins across cell types

To investigate the cell-specific profiles of altered RG4BPs between normal and AD individuals, seven single-cell datasets containing 59 normal and 61 AD individuals were collected (**Figure 1a, Supplemental Table 1**). In addition to the 1100 RG4BPs documented in the QUADRatlas ^27^, 318 of which are supported by experimental evidence, we included 130 HTR-SELEX tested RG4BPs (37 are newly identified) that exhibit strong rG4 enrichment ^28^(**Supplemental Table 2**). The integrated dataset underwent a quality control (QC) prior to batch correction, resulting in the removal of 19% of low-quality cells and potential doublets, with 601,213 cells retained for downstream analysis. After batch correction and dimensionality reduction, six major clusters were annotated using corresponding known markers (**Figure 1b**, see **Methods**). The cross-validation using identity pre-labelled datasets demonstrated a high concordance, with 98% accuracy and 99% recall (**Supplemental Fig. 1**). Pericytes and endothelial cells were not individually annotated, as previous datasets lacked distinguishable marker genes and were therefore not used for subsequent analyses. With major cell clusters annotated, a total of 149 differentially expressed RG4BPs between AD and normal individuals were identified using previously applied criteria to detect subtle alteration ^29^ (**Figure 1c**, see **Methods**).

**Figure 1.**
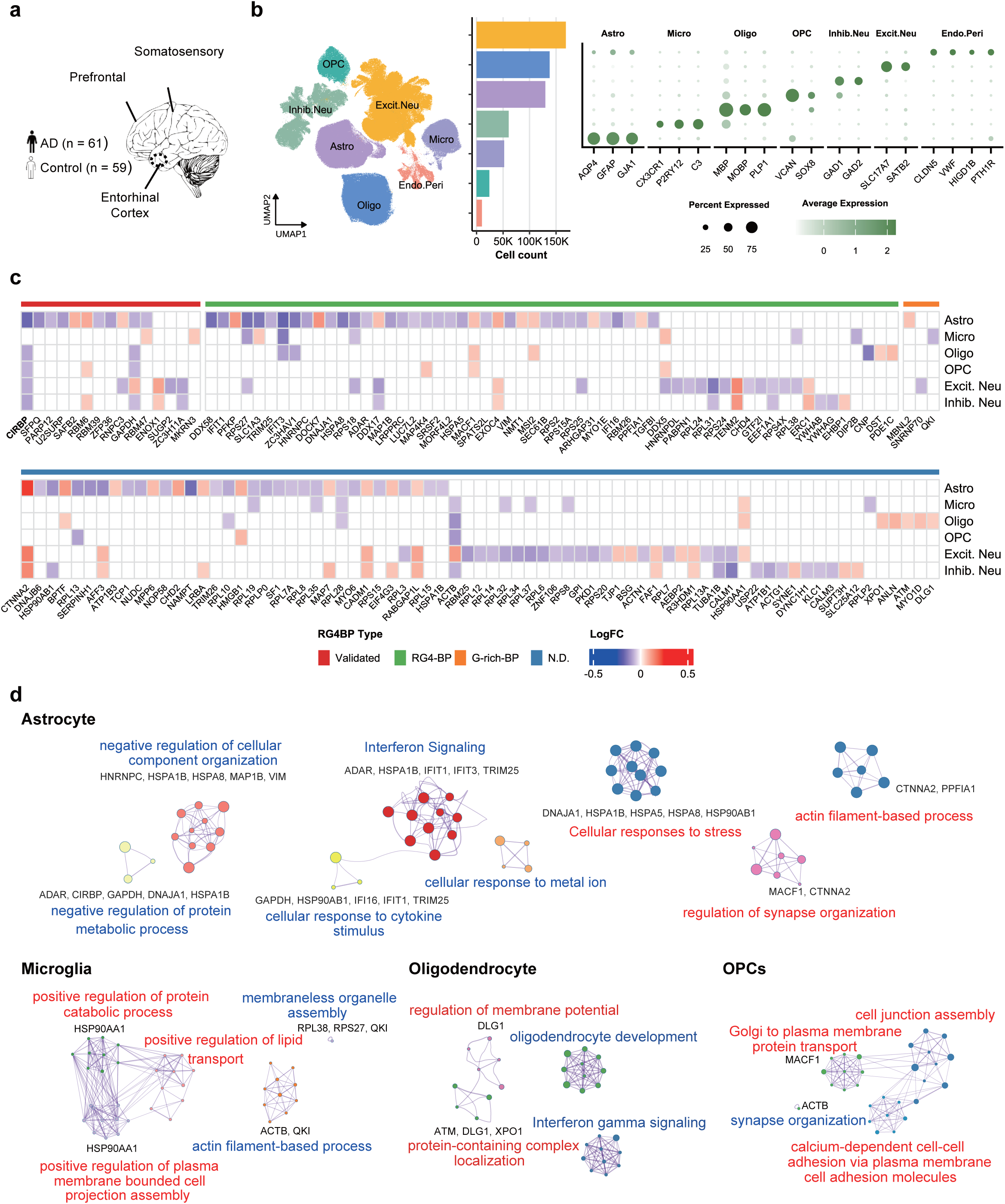
Integrated snRNA-seq analysis and rG4-binding protein expression landscape. (a) Schematic overview of sampled brain regions (PFC, somatosensory cortex, and entorhinal cortex) and cohort composition (AD, n = 61; control, n = 59) across seven integrated snRNA-seq datasets. (b) Re-annotated seven major brain cell types. (c) The log fold change of differentially expressed RNA G-quadruplex binding proteins (rG4BPs) between AD and normal individuals across major cell types. RG4BPs were grouped into four classes (1) RG4-BPs, (2) G-rich-BPs, and (3) proteins with undetermined binding status (N.D.), all derived from the QUADRatlas database, and (4) validated RG4BPs, defined as those experimentally supported by HTR-SELEX. (d) Functional enrichment network of differentially expressed genes (DEGs) between AD and control groups. Blue clusters represent downregulated pathways, while red clusters represent upregulated pathways. Genes highlighted within each cluster denote RG4BPs.

Remarkably, several known RG4BPs catalogued by QUADRatlas^27^, including CIRBP, HSP90AA1, and SLC1A3, were overtly dysregulated, given the fact that many RG4BPs were distinctly expressed across cell types (**Supplemental Table 3**). Functional enrichment analysis of differentially expressed genes revealed that AD-dysregulated RG4BPs were involved in diverse lineage-specific cellular functions. Compared to non-AD individuals, many downregulated RG4BPs in astrocytes were enriched in Interferon signaling, cellular responses to stress, cellular response to cytokine stimulus, negative regulation of protein metabolic process, and negative regulation of cellular component organization. Upregulated RG4BPs in astrocytes were mainly enriched in actin filament-based processes, response to metal ions, and regulation of synapse organization. In microglia, dysregulated RG4BPs were enriched in actin filament-based process, membrane-less organelle assembly, positive regulation of protein catabolic process, positive regulation of lipid transport, and positive regulation of plasma membrane bounded cell projection assembly. While in oligodendrocytes, dysregulated RG4BPs contribute to protein-containing complex localization, interferon gamma signaling, oligodendrocyte development, regulation of membrane potential, and regulation of postsynaptic membrane neurotransmitter receptor levels, expression altered RG4BPs in OPC were primarily enriched in cell junction assembly, golgi to plasma membrane protein transport, calcium-dependent cell-cell adhesion via plasma membrane cell adhesion molecules, and synapse organization (**Figure 1d**). These expression-altered rG4BPs in oligodendrocytes (OLG) and oligodendrocyte progenitor cells (OPCs) potentially contribute to the impaired cell communications, myelination, and neuroinflammation of AD patients ^30,31^. Of note, among these RG4BPs, the stress-sensor cold-induced RNA-binding protein (CIRBP) is systematically downregulated across types, suggesting its important role in AD-specific dysregulation.

### Cell type-specific response of RG4BP target genes

The gene expression of CIRBP target genes was then examined across distinct cell types, as CIRBP exhibits both protective and cytotoxic roles in AD ^32,33^. Experimentally profiled binding sites of CIRBP in human and multiple datasets obtained in mouse were compiled into a non-redundant target gene list regardless of the cell identity based on orthologs (**Figure 2a**), with the rationale that CIRBP and many genes are evolutionarily conserved between human and mouse ^34^, and RNAs of target genes are accessible when expressed. This resulted in 1459 candidate target genes, of which 208 were differentially expressed between AD and normal individuals (**Figure 2b**). Although CIRBP has been previously reported to bind linear sequences, we identified 1388 expressed CIRBP target genes (192 differentially expressed) harbouring loci exhibiting strong rG4-forming potential (**Supplemental Table 4**) in the human transcriptome. Intersecting with experimentally validated and predicted binding sites in QUADRatlas, at least 1409 candidate target genes contain validated or predicted rG4-forming sequences, suggesting that binding to rG4 is a part of CIRBP’s biological roles. Indeed, our HTR-SELEX experiment confirmed the enrichment of rG4, and the validated rG4 binding was confirmed using EMSA (**Supplemental Fig. 2**). Notably, distinct cell types exhibited lineage-specific CIRBP binding targets.

**Figure 2.**
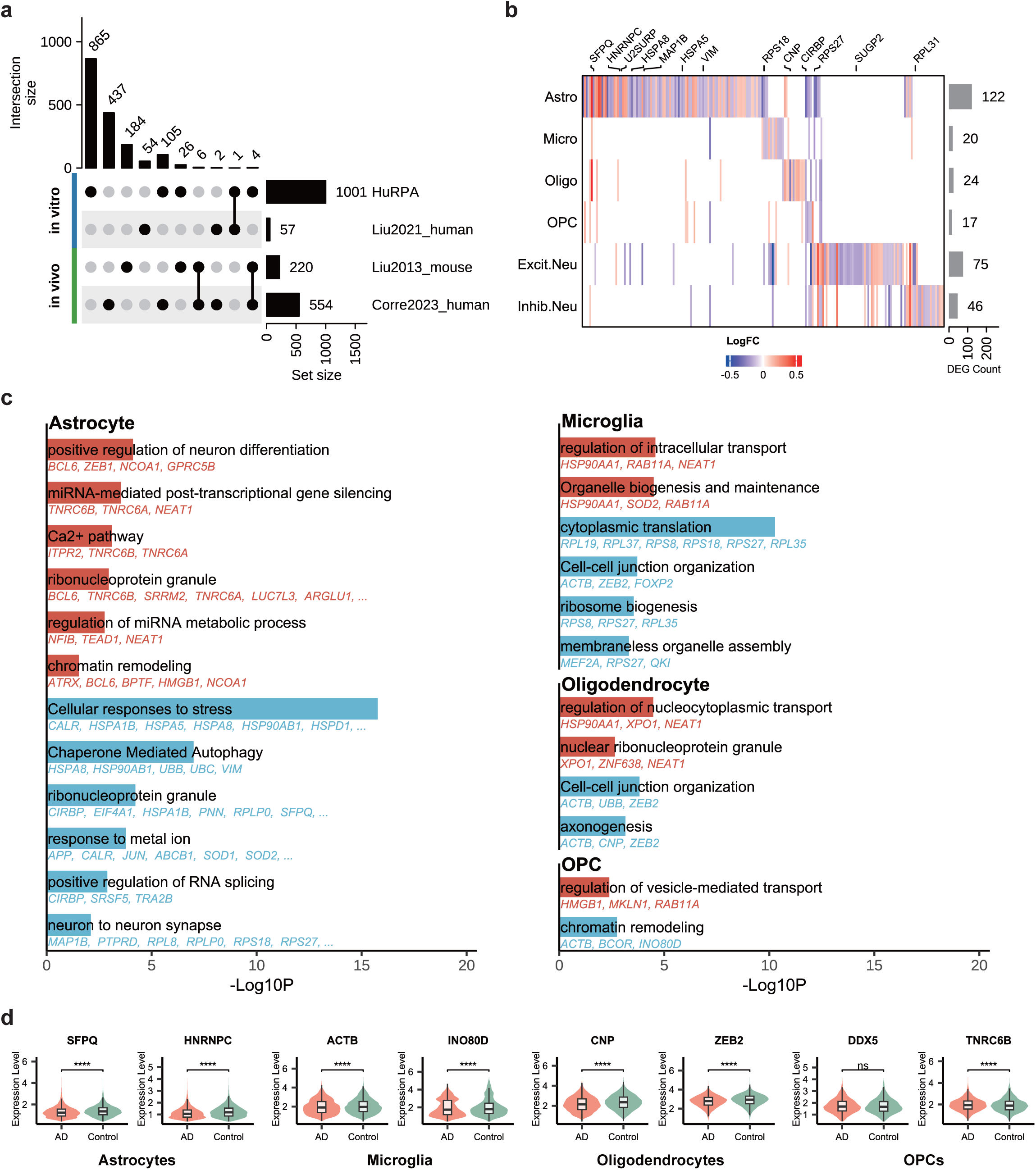
Altered expression and functional roles of CIRBP-regulated target RNAs in AD. (a) The intersection of CIRBP targets derived from four sources and their respective set sizes. CIRBP target genes were collected from two *in vivo* and two *in vitro* experiments, and target genes identified in the mouse transcriptome were converted to human gene IDs through biomaRt. (b) The log fold change of differentially expressed CIRBP target genes across major brain cell types between AD and control individuals. (c) Enriched Biological Process (BP) GO terms of CIRBP target genes. Blue and red represent enriched functional terms derived from down-regulated and up-regulated genes, respectively. Representative top five genes within each term were labelled under each corresponding bar. (d) Known and newly detected genes that are significantly dysregulated across four key brain cell types, presented by violin plots using normalized expression levels. Two-side Wilcoxon rank-sum test between AD and control groups (symbols indicate significance levels: * p < 0.05, ** p < 0.01, *** p < 0.001, **** p < 0.0001, ns not significant).

Next, functional enrichment was executed to gain further biological insights into CIRBP target genes. Several significantly enriched biological processes are directly linked to AD pathological features (**Figure 2b**). Considering that rG4 directly participates in the cellular stress response, such as oxidative stress, hypoxia, and DNA damage, and CIRBP is a stress-response gene that shuttles between the nucleus and cytoplasm ^35^, we reason that dysregulated cell stress response contributes to chronically impaired cellular function. Distinct cell types present diverse outcomes as they cooperatively play distinct biological roles in maintaining systemic homeostasis. Indeed, the dysregulated CIRBP target genes in astrocytes were functionally enriched in the response to metal ion, cellular responses to stress, ribonucleoprotein granule (both down and up), neuron to neuron synapse, chaperone mediated autophagy, positive regulation of RNA splicing, chromatin remodeling (up), positive regulation of neuron differentiation (up), regulation of miRNA metabolic process (up), Ca^2+^ pathway (up), and miRNA-mediated post-transcriptional gene silencing (up). In microglia, enriched biological processes with RG4BPs included intracellular transport, organelle biogenesis and maintenance, ribosome biogenesis, cytoplasmic translation (down), cell-cell junction organization (down), and membraneless organelle assembly (down). Differentially expressed genes in oligodendrocytes were enriched within cell-cell junction organization (down), axonogenesis (down), regulation of nucleocytoplasmic transport, and nuclear ribonucleoprotein granule. In contrast, differentially expressed CIRBP target genes were enriched in chromatin remodeling (down) and regulation of vesicle-mediated transport within OPC **(Figure 2c**). These observations align with the physiological identity of corresponding cells in the CNS.

This strategy identified dysregulated known genes in astrocytes in the AD context, including SFPQ, HNRNPC, APP, NOCA1, and BCL6, in addition to many ribosomal proteins and elongation factors, all of which are targeted by the CIRBP protein. Besides transcripts of DDX5, TNRC6B, and RAB11A, which are captured as CIRBP protein targets in OPCs, the ZNF638 RNA, which is responsible for repairing in multiple sclerosis, was found to be involved in chromatin remodeling ^36^. Two downregulated genes, ACTB and INO80D, which can form a complex to modulate epigenetic landscapes, were also identified as CIRBP targets ^37^. In oligodendrocytes, CNP and ZEB2, which respectively regulate axonal degeneration and influence myelin integrity, were downregulated ^38,39^(**Figure 2d**). The stress-related genes, HSP90AA1 and *Neat1*, were upregulated. Unexpectedly, many CIRBP target RNAs also encode RG4BPs, implying an inter-tangled rG4-associated regulation. Indeed, many differentially expressed genes targeted by RG4BPs contain PQSs that are also associated with various RNA modifications ^40^ (**Supplemental Fig. 3**). Collectively, impaired functions of these CIRBP target genes demonstrate that RG4BPs regulate essential cell-specific processes to maintain their physiological characteristics and become functionally dysregulated in AD pathology (**Figure 2c**).

### RG4BPs in glial cells distinctly impact transcriptional states

Given that astrocytes and microglia maintain brain homeostasis, and both cell populations exhibit heterogeneous transcriptional states, we set out to investigate the functional impacts of dysregulated RG4BPs on AD etiology and progression. As the transcriptional states in the aforementioned combined datasets from multiple sources could not be clearly resolved, an independent astrocyte dataset containing five brain regions from 32 individuals was employed instead, considering its design to mitigate batch effects ^12^. Upon successful sub-clustering of distinct transcriptional states based on comprehensive annotation by integrating Jaccard index and known markers (**Figure 3a**, **Supplemental Fig. 4**, and **Supplemental Table 5**), we examined differentially expressed RG4BPs and CIRBP target genes in each sub-cluster (**Figure 3b** and **Supplemental Fig. 5a**). In line with the previous population level observation, the expression of RG4BPs (67% of previously detected RG4BPs were detected here) and CIRBP target genes was significantly altered across sub-clusters, indicating cells at distinct transcriptional states were asymmetrically impacted. Although the direction of some genes changed counterintuitively, this could be explained by subtle alterations within a single state and by the effect of addiction on aggregation (**Supplemental Fig. 5b**). Thus, we further investigated state-specific changes of RG4BPs in astrocytes exhibiting homeostatic (astH0), reactive (astR1 and astR2), intermediate (astIM), and metabolic (astMet) states, without considering potential doublets astNeu and astMic (see **Methods**). Within the differentially expressed genes between normal and AD individuals across transcriptional states, approximately 33% to 47% genes expressed RNAs targeted by RG4BPs, while only about 7 % of differentially expressed genes are RG4BPs (**Supplemental Fig. 5c**). Dysregulated RG4BP (HSPA1A and HSPA1B) were enriched in positive regulation of supramolecular fiber organization and protein refolding in astH0 (HSPA1A, HSPA1B, HSP90AA1, HSPB1, CRYAB), modulation of chemical synaptic transmission in astR1 (up), distinct set of genes of which were also downregulated in reactive astrocytes astR1 (**Figure 3c and Supplemental Table 7**). Several RG4BPs in the response to oxidative stress (HSPA1A, HSPA1B; GO:0006979) were significantly upregulated in astH0 and astR1. This indicated that an altered stress response is associated with the transition of astrocyte states. However, the expression profiles of stress-granule related genes were not significantly changed (**Supplemental Fig. 6**), suggesting stress granule independent stress response in astrocytes ^41^.

**Figure 3.**
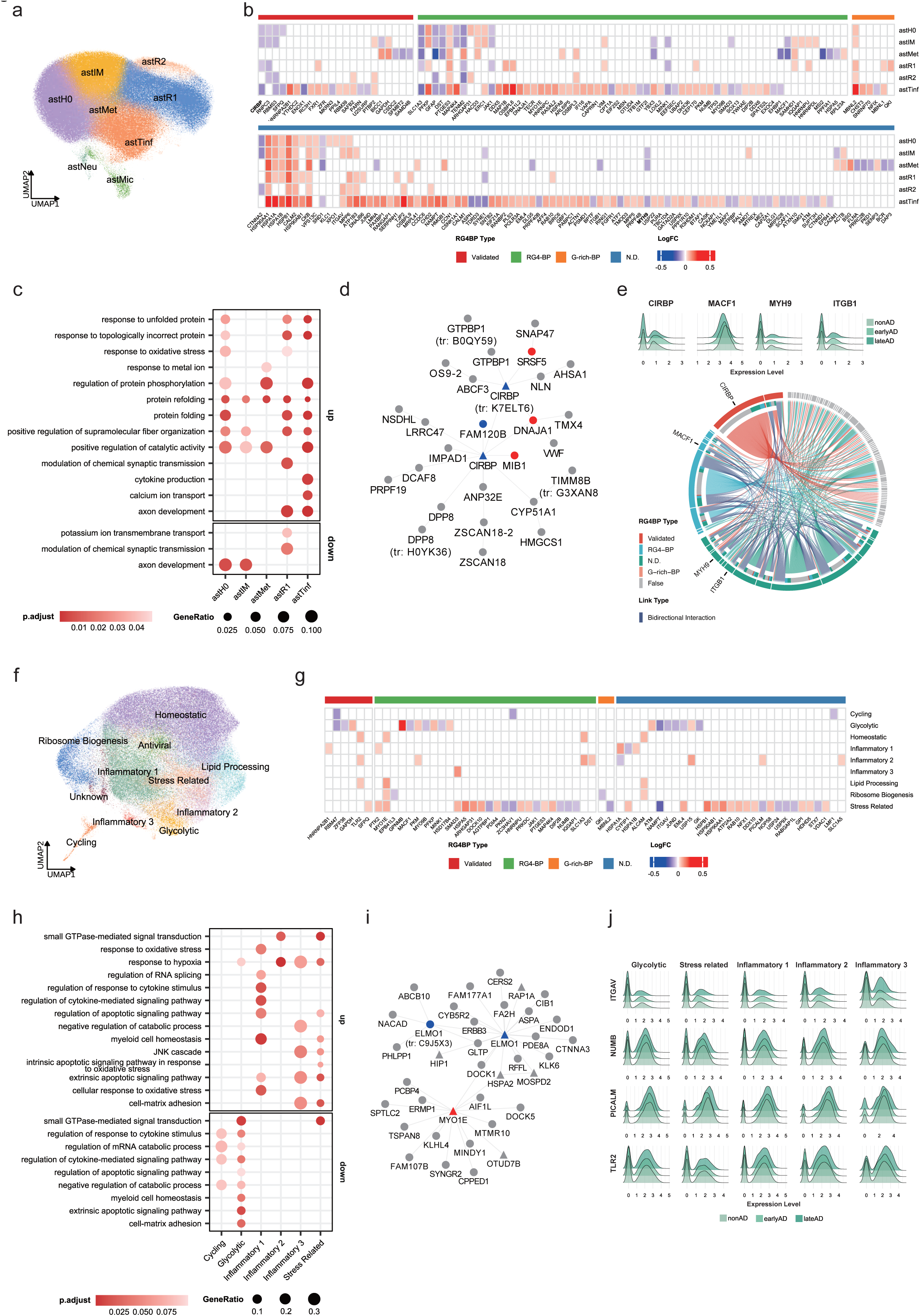
Transcriptional heterogeneity of astrocyte and microglia associated with AD. **Astrocytes (a–e):** (a) Astrocyte subcluster identification and annotation. (b) The log fold change (logFC) of four rG4BP categories (Validated, RG4-BP, G-rich-BP, and N.D.) across astrocyte subclusters. (c) Representative Biological Process (BP) GO terms enriched in each astrocyte transcriptional state, highlighting the functional roles of RG4BPs. (d) Bayesian causal network of the key driver protein (KDP) CIRBP. Node shapes indicate classification: triangles and circles represent KDPs and non-KDPs, respectively. Node colors denote differential gene expression in astMet compared to earlyAD and lateAD: upregulated, downregulated, and non-significant or unexpressed genes are labelled in red, blue, and gray, respectively. The accession numbers of proteins from TrEMBL are indicated. Protein isoforms are indicated with a suffix. (e) The relationship between CIRBP and its target genes. Outer and inner rings indicate the rG4BP type of the genes and their linked partners, respectively. Arrows denote the targeting direction, with bidirectional bindings shown in dark blue. Three key targets (MACF1, MYH9, and ITGB1) are highlighted. Accompanying ridge plots demonstrate the normalized expression levels of these genes in the astMet subcluster throughout AD progression (nonAD to lateAD). **Microglia (f–j):** (f) Microglia subcluster identification and annotation. (g) The logFC of RG4BP across microglia subclusters. (h) Representative BP GO terms of RG4BPs across microglia transcriptional states (i) Bayesian causal network of the KDP MYO1E. (j) The normalized expression levels of four key genes (ITGAV, NUMB, PICALM, and TLR2) across five distinct microglial subclusters throughout AD progression.

The dysregulated supramolecular fiber organization in both astH0 and reactive astrocytes further restates the importance of RG4BPs associated with non-membrane granule assembly. However, distinct sets of genes were upregulated in homeostatic astrocytes and reactive astrocytes (**Figure 3c**). Interestingly, the down-regulated cellular response to potassium ion and potassium ion transport in reactive astrocytes manifested an impaired potassium homeostasis (SLC1A3), potentially influencing the normal binding between RG4BPs and rG4s. Worth mentioning, CIRBP was previously identified as a key driver protein, highlighting its essential role in AD (**Figure 3d**) ^25^. Several RG4BPs (DST, FLNA, ITGB1, MACF1, and MYH9), involved in impaired wound healing, were downregulated in astMet. While CIRBP directly targets ITGB1, MACF1, and MYH9 RNA, CIRBP RNA can interact with ITGB1, FLNA, and MYH9 protein. The coordinated downregulation of paired genes implies their direct regulation and contribution to the malfunctioning signalling pathways (**Figure 3e**). This was accompanied by upregulated synapse organization (including APP and APOE) in astR1, in which RG4BP MAP1B directly regulates cytoskeleton remodelling, as does FLNA (**Figure 3e)**. Although CIRBP protein exhibits an age-associated increase, the temporal expression of CIRBP shifted from up-regulation in early AD to down-regulation in late AD, indicating an altered regulatory mechanism of the stress response during AD progression, which can explain the dual roles of CIRBP (**Supplemental Fig. 7**). Upon checking RG4BP target RNAs in astrocytes, we found that 44 % and 47% of them were differentially expressed between AD and control and between early-AD and control, respectively. At least 95% of these target RNAs contain rG4-forming loci in their longest transcripts, suggesting an undisclosed relationship between rG4 and AD etiology. Moreover, about 63% of these differentially expressed RG4BP target genes were upregulated, suggesting they partially contribute to the accumulated rG4s (**Supplemental Fig. 8**).

Like astrocytes, an independent microglia dataset was examined to investigate dysregulated RG4BPs across transcriptional states (**Figure 3f**, **Supplemental Fig. 9, and Supplemental Table 6)**. As expected, suppressed chemotaxis-related processes correlate with microglia activation in AD ^42^. SMAD3, which contributes to microglial activation by regulating cell junction assembly (GO: 1901888), was significantly upregulated. This corresponds to its impaired crosstalk with astrocytes, as SMAD3 was downregulated considerably in astIM and astTinf. In the stress-related microglia, five RG4BPs, including SMAD3, SNCA, DOCK10, PTK2, and MAP4K4, were significantly upregulated, underscoring the important functional roles of dysregulated RG4BPs ^43^ (**Figure 3g**). Despite the upregulation of microtube-remodel gene MYO1E across distinct states, several key RG4BPs (ITGAV, PICALM, TLR2, and NUMB) regulating endocytosis were dysregulated in glycolytic, stress-related, and inflammatory microglia. TLR2 and PICALM were, however, upregulated in inflammatory microglia, while PICALM and other genes were downregulated in glycolytic, stress-related microglia (GO:0030100) (**Figure 3g**). Of note, the PICALM isoform has been identified as medically relevant ^44,45^, which can cause abnormal lipid droplets during endocytosis due to compromised lysosomal function ^46^. Meanwhile, we also found that dysregulated RG4BP could reshape the transcriptional state, as some genes in stress-related microglia (HIF1A, OXR1, LRRK2, FOXO3, FOXP1, PRKN, TP53INP1, FOXO1, KAT2B, SLC8A1, and HSPA1A) were enriched in the “cellular response to oxidative stress”. While these genes in the AD group exhibited significantly downregulated expression, a distinct set of genes (ETV5, ATP2A2, NCOA7, MAP3K5, SNCA, SIRPA, HSPA1B) were upregulated (**Figure 3h** and **Supplemental Fig. 10**). Such complex expression changes indicate a complex response that is potentially related to AD progression. Indeed, the dysregulated MYO1E and ELMO1 encode key driver proteins in AD (**Figure 3i**), highlighting their fundamental roles in impaired endocytosis. By examining the differences between differentially expressed genes derived from early and late AD stages (**Figure 3j and Supplemental Table 7**), the dysregulated genes were indeed associated with initial stress, apoptotic signaling, and neurotoxicity induced by persistent stress (**Figure 3g** and **Supplemental Fig. 11**). Taken together, malfunctioned transitions of transcriptional states in astrocytes and microglia directly impaired key RG4BPs associated with AD hallmarks.

### RG4BPs participate in essential state-specific regulatory processes

To elucidate state-specific RG4BP regulators, gene modules across transcriptional states were constructed using highly variable genes combined with expressed RG4BP and RG4BP targets, which were subjected to network and pathway analyses (see **Methods**). Eight and eleven modules were identified in astrocytes and microglia, respectively (**Figure 4a, b**). Many expressed RG4BPs in astrocytes are highly representative in modules AST-M1, AST-M2, and AST-M4. Whereas a distinct set of RG4BPs is associated with modules MIC-M2, MIC-M4, and MIC-M8 in microglia (**Figure 4a and Supplemental Table 8**).

**Figure 4.**
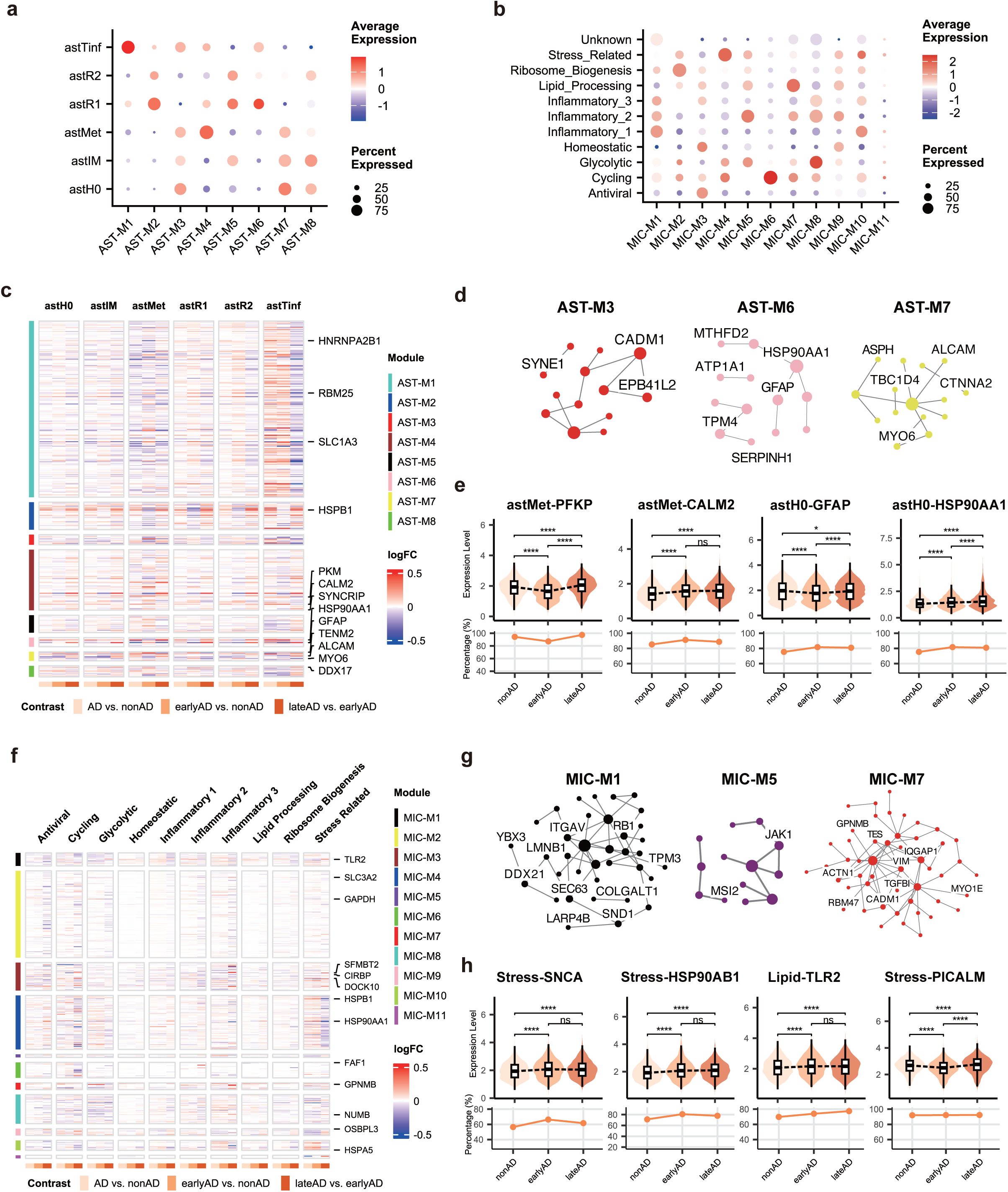
Integrated co-expression and PPI networks uncover AD-specific RG4BP dynamics in glial cells. (a, b) State-specific module characteristics in astrocyte (a) and microglia (b) transcriptional states. (c, f) Differential expression profiles (LogFC) of module genes across AD progression (non-AD to late-AD) in astrocyte (c) and microglia (f) subclusters. (d, g) Integrated co-expression and PPI networks for representative modules in astrocytes (d) and microglia (g). Labeled nodes highlight key RG4BPs and representative genes with distinct trajectory patterns across disease states. (e, h) Temporal expression dynamics of AD-dysregulated RG4BPs and corresponding patterns across AD stages in astrocyte (e) and microglia (h) subclusters.

In astrocytes, more than 120 RG4BPs (**Figure 4c**), annotated by either QUADRatlas or HTR-SELEX, were found in gene modules, and several known AD-dysregulated genes, including PFKP, HSP90AA1, GFAP, SERPINH1, CADM1, and SYNE1, were either co-expression or protein-protein interaction hubs (**Figure 4d**). For instance, PFKP that regulates canonical glycolysis, was dysregulated at the early stage of AD and became overtly upregulated over time, suggesting its transcriptional state shift toward exhaustion (**Figure 4e** and **Supplemental Fig. 12**). This is consistent with the characteristics of module 4, which regulates cellular response to hypoxia, pyruvate metabolic process, and intracellular monoatomic cation homeostasis (**Supplemental Fig. 13**). Indeed, genes controlling ion homeostasis were dysregulated in AD patients, consistent with existing reports on ion imbalance, with many of the affected genes regulating zinc, copper, and calcium homeostasis ^47^. In particular, RG4BP calmodulin 2 (CALM2) was consistently upregulated in early and late stages of AD patients. The CALM2 protein potentially coordinated with dysregulated metallothionein proteins at the early stage of AD to exacerbate imbalanced ions. RG4BPs in module 7 are significantly enriched in axon guidance (EPHA4; ALCAM, EPHA6, NRXN1, EFNA5, and EPHB1), cell-cell adhesion via plasma-membrane adhesion molecules (ALCAM, CADM2, NRXN1, LRRC4C), and supramolecular fiber organization (COL5A3, MYO6, and SHROOM3). In module 6, the known astrocyte marker GFAP was identified as a hub, enriched for supramolecular fiber organization. However, GFAP was downregulated in the early stage and became upregulated in the late stage, consistent with the reported shift in its regulatory role ^48^. Worth mentioning, the GFAP interacting HSP90AA1 (**Figure 4d**) was upregulated over time during the progression of AD, except for reactive astrocytes and astMet, which was further supported by the proportion of cells expressing the corresponding gene (**Figure 4e** and **Supplemental Fig. 12**). Because chaperone HSP90AA1 regulates unfolded protein, the impaired interaction between HSP90AA1 and GFAP potentially contributes to the exacerbated cellular stress response. In reactive astrocytes, HSP90AA1 was downregulated at the early stage, but upregulated in astH0 and astTinf, without alteration in astMet and astIM. Counterintuitively, HSP90AA1 became overtly upregulated in the late AD stage in stress-related astrocytes astMet. The dynamics of these RG4BPs revealed a shift from the acute stress response to a chronic stress state. Importantly, dysregulated RG4BPs were targeted by astrocyte-specific transcription factors. Inferred regulation was found between RFX4 and GFAP, VIM, PLEC, MACF1, and PKFP, GAFP in PFC, while MACF1 and SLC1A3 were regulated by EMX2 ^24^. In contrast, SAFB2 and PLEC were regulated by RFX2 in EC. Notably, VIM, GFAP, and PLEC were identified as putative key driver proteins in AD ^25^, emphasizing that RG4BPs are integral to gene regulation.

As microglia gradually lost their ability to efficiently clean Aβ plaques, the differentially expressed RG4BP associated with AD etiology (AD vs. control) and progression (early vs. late) were examined separately (**Figure 4f**). The representative functional enrichment of each module corresponds to the biological characteristics of each transcriptional state, confirming the state-specific dysregulation (**Supplemental Fig. 13**). Many (168) RG4BPs were identified as hubs in protein-protein interaction networks, particularly those responsible for binding unfolded protein, which became dysregulated in stress-related microglia (**Figure 4g**). For instance, the central oxidative stress regulator HIF1A, which also regulates chemokine−mediated signaling pathways and cellular response to oxidative stress in glycolytic microglia, became dysregulated together with RG4BPs (GO:2001233) (HIF1A, ITGAV, and BCL2) (**Supplemental Fig. 14)**. Additionally, the RNA level of SNCA was upregulated in microglia (**Figure 4h**). Given that SNCA is a known RG4BP ^49^ and its transcripts are targets of RG4BPs ^26^, we hypothesized that some RG4BPs were dysregulated. Consistent with this, HSP90AB1 that targets SNCA transcripts was found to be upregulated in glycolytic and stress-related microglia at the early stage of AD, reflecting a transcriptional shift toward activation and stress response. The stressed-related microglia exhibit significant changes toward protein folding (HSPB1, HSP90AB, HSP90AA1, CHORDC1, HSPD1, BAG3, PTGES3, HSPH1, ST13, HSPA4L) and regulation of protein stability (HSP90AB1, HSP90AA1, SNCA, HSPD1, BAG3, PTGES3). While overall expression of chemokine production genes (HIF1A, CXCR4, TLR2, PTK2B) was downregulated in AD, the late AD exhibited a slight upregulation of HIF1A, suggesting intensified hypoxia response in late AD that potentially causes pro-inflammatory bias and transit cells into a more glycolysis state, therefore worsening the neuroinflammation and reducing the Aβ clearance efficiency. Indeed, genes (PRKAG2, DDIT4, HDAC4, HIF1A, INSR, and KAT2B) in the regulation of glycolytic process (GO:0006110) were upregulated in stress-related microglia in late AD compared to the early AD. In glycolytic microglia, genes involved in the response to lipopolysaccharide (SLC11A1, SLC7A5, IRAK2, ZFP36, TNFRSF1B, PDE4B) and regulation of the response to cytokine stimulus (PTPN1, IRAK2, PADI2, HIF1A) were downregulated, further supporting the observation in stress-related microglia. By contrast, in lipid-processing microglia, dysregulated TLR2 was associated with regulation of the membrane raft (**Figure 4h**). We further examined the expression of constituent proteins of microglial extracellular vesicles and found that APP was only slightly downregulated in AD, but not between early and late stages. The overtly upregulated HSP90AA1, however, exhibited a decrease at the late stage compared to the early stage. This potentially reflects a chronic stress-induced functional exhaustion, as SOD1 only became subtly downregulated in late stages ^50,51^. These altered genes markedly contribute to chronic stress and the pro-inflammatory response of microglia, together with compromised lysosomal function mediated by PICALM, resulting in insufficient Aβ clearance and phagocytic exhaustion (**Figure 4h**).

These findings demonstrate that RG4BPs exert state-specific and essential regulatory roles in maintaining homeostasis under normal physiological conditions and become dysregulated upon the onset of AD, even though rG4 structure is not chronically accumulated in microglia ^22^.

## Discussion

Non-canonical RNA secondary structure rG4 and its interacting RG4BPs are promising but understudied players in AD pathogenesis ^18–22^. Their cell-type-specific and state-resolved functions remain poorly characterized. We integrated multiple single-cell transcriptomic profiles to delineate RG4BP dysregulation across cell types and transcriptional states, uncovering central roles of RG4BPs in existing AD hypotheses.

The lineage-specific alterations of RG4BP expression suggest that they regulate essential cellular functions related to cell identity, including ion homeostasis and synapse organization in astrocytes, protein catabolic process and lipid transport in microglia, and regulation of membrane potential in oligodendrocytes. In line with previous reports on the accumulation of RG4 in astrocytes ^17,20,21^, the many RG4BPs in astrocytes were downregulated in AD, including CIRBP, SFPQ, BSG, SLC1A3, and ENOX1. Their overall downregulation at the cell-type level highlights astrocytes as a primary cellular context in which rG4–RG4BP dysregulation may exert functional consequences. In the meantime, more than 50% of upregulated genes contain rG4-forming loci in their longest transcripts, suggesting an impaired mechanism for resolving rG4 *in vivo*. The stress-sensor CIRBP was systematically downregulated across all cell types, aligning with its dual roles in AD and suggesting a broadly compromised, RG4BP-mediated stress response. Functional enrichment of CIRBP target genes reinforces this speculation, as dysregulated target genes in astrocytes are enriched in pathways for chaperone-mediated autophagy, Ca²⁺ homeostasis, and ribonucleoprotein granule assembly, while targets in microglia are enriched in pathways for organelle biogenesis and cell-cell junctions. These processes are lineage-specific and critical for maintaining brain homeostasis. Remarkably, more than 90% of the CIRBP target genes contained rG4-forming loci with a relatively conservative estimation, indicating a previously underappreciated non-random association between rG4 and AD.

The altered RG4BP expression in distinct states, however, counterintuitively differs from the average signal of the entire cell population, exhibiting state-specific upregulation. The upregulated RG4BPs in astH0 in early-state AD patients are involved in regulating protein refolding and stress granule assembly, reflecting an early acute stress response. In contrast, downregulated genes in reactive astrocytes (astR1/astR2) and astMet are mainly involved in the regulation of stress response (CIRBP and YTHDF1) and impaired potassium ion transport. These findings support the observation that the transcriptomic profiles of astMet in late AD patients are distinct from those in early AD patients ^12^, with a more pronounced impairment of ion homeostasis. Such changes align with recent observations that Li^+^ is trapped or sequestered by Aβ plaque ^23^, disrupting cation homeostasis, as Li^+^ destabilizes rG4. This convergence supports a model in which chronic ion imbalance promotes rG4 accumulation during late-stage AD. The temporal shift in CIRBP from early upregulation to late downregulation further supports a model in which acute stress responses evolve into chronic astrocyte exhaustion, driven by sustained RG4BP dysregulation and maladaptive transcriptional state transitions ^52^. This shift could be mediated by or lead to progressive rG4 accumulation in late AD, which impairs intracellular shuttling of CIRBP and thereby worsens the stress response ^35^.

Dysregulated RG4BPs in microglia impair microglia activation and stress response. In addition to cytoskeleton-associated genes, many dysregulated RG4BPs affect phagocytosis, protein folding, and immune response in the stress-related subcluster during the onset of AD. These alterations suggest that rG4–RG4BP dysregulation impairs microglial stress sensing and functional responsiveness, thereby influencing transcriptional state transitions. Indeed, an overall dysregulation of stress-related biological processes was observed by the upregulation of genes in receptor-mediated endocytosis and the downregulation of phagocytic activity in stress-responsive microglia. Notably, while the HIF1A-associated chemokine response was suppressed in early AD stages, it became upregulated in late-stage AD, indicating intensified oxidative stress as the disease progresses ^53^. Meanwhile, stressed microglia exhibit a reduced PICALM expression, leading to accumulated lipid droplets that impair endocytosis and thereby hijack normal Aβ clearance (downregulated in glycolytic and stress-related microglia). Although PICALM was upregulated in inflammation 2 microglia, corresponding lysosomal components were downregulated, suggesting keeping specific isoforms is central to microglial function^46^.

Collectively, our findings outline dysregulated RG4BPs as a potential molecular link between stress responses, ion imbalance, and glial dysfunction, positioning rG4–RG4BP interaction as a mechanistic layer that integrates transcriptional state failure with chronic glial stress in AD. We provide a plausible mechanistic basis for connecting discrete pathological features that might converge to drive AD through dysregulated RBP-RNA interactions. However, the current investigation is limited to single-cell transcriptomic profiles and does not account for full RNA transcripts, proteome, or RNA modifications. In addition to lacking full transcript information for the protein-binding site analysis, changes in RNA levels do not necessarily alter protein or RNA modifications, even though key enzymes, like YTHDF1 and ADAR, are dysregulated. Further mechanistic studies shall be conducted to confirm these findings and provide an in-depth analysis of RNA isoforms and corresponding RG4BP binding sites in primary glial cells using *in vivo* profiling assays, such as eCLIP, SPIDR, and MPIT ^54,55^. Nevertheless, our study explored the interactome between RBPs and rG4 at the cell-type and transcriptional-state resolutions, underscoring the importance of examining the interactions between RBPs and RNA structures in neurodegenerative diseases.

Many RBPs, including SNCA, HSP90AA1, PICALM, HNRNP1, and TDP-43, are not only directly associated with rG4, but also play essential roles in other neurodegenerative diseases. Given the shared functional pathways across distinct neurodegenerative diseases ^56^, it is reasonable to speculate that RG4BPs play a broader role. Although the precise functional roles of RG4BPs remain to be disclosed, understanding their dysregulated interactions with RNA secondary structures in the CNS provides novel insights into molecular mechanisms and open an alternative avenue for designing RNA structure-based therapeutic strategies.

## Methods

### RG4BP targets and CIRBP target supported by experimental evidence

The list of RG4BPs catalogued in QUADRatlas was combined with HTR-SELEX detected RBPs showing high rG4 enrichment ^28^. RG4BP targets were primarily obtained by merging immunoprecipitation datasets and experimentally identified interactions reported in HuRPA (https://sysbiocomp.ucsd.edu/prim/) ^26^.

Three public datasets (GSE164523, GSE40468, and GSE225516) were analyzed to obtain human CIRBP target transcripts. For the SELEX of human CIRBP (GSE164523^57^), peaks were assigned to genes of human hg19 gene annotation (UCSC Feb. 2009 GRCh37/hg19) using the intersect function of bedtools by including at least 50% region of the original peaks in the gene. Corresponding gene symbols were converted from Ensembl Gene IDs using biomaRt package (2.60.1, Ensembl release 75). Genes supported by at least one CIRBP peak were defined as candidate CIRBP targets. For the CIRBP RIP-seq data (GSE225516^58^) derived from the human LoVo colon cancer cell line, RIP-seq data under infected and non-infected conditions generated by CIRBP immunoprecipitation (CIRBP IP), IgG control RIP, and input total RNA were downloaded, of which there are at least two replicates for each experiment. raw sequencing data were mapped to the GRCh38 genome using Bowtie2 v2.5.4. Samtools v1.21 and featureCounts v2.0.8 were used to sort bam files and perform reads counting. CIRBP-specific enrichment over background was performed separately for infected and non-infected conditions with DESeq2 v1.44.0. For each condition, differentially enriched genes were defined as those with padj < 0.05 and log2FC ≥ 0.5 in CIRBP vs Input group, as well as padj < 0.05 and logFC ≥ 0.25 in CIRBP vs IgG group, the intersection of these two gene sets per condition, and the combination across two conditions were considered as candidate CIRBP pull-down targets. For the CIRBP PAR-CLIP dataset derived from mouse embryonic fibroblasts (GSM994702^59^), a similar strategy was applied to the mouse mm9 genome (UCSC Jul. 2007 NCBI37/mm9) with the same bedtools intersect parameter to get Ensembl gene IDs. Mouse gene IDs were then converted into corresponding human orthologs using biomaRt (version 2.60.1, Ensembl release 114). Dataset-specifically and overlapping gene lists were then obtained.

### Genes containing rG4/PQS are associated with RNA modifications

The longest transcripts of human genes were used to predict rG4 forming loci, followed a hierarchical prediction to obtain high reliability loci^40^. RG4BPs target genes of was counted as rG4-containing if at least one rG4-forming locus was predicted. The rG4-associated RNA modifications were extracted from the MoRNiNG database.

### snRNA-seq data processing and cell type annotation

Several public single-cell or single-nucleus RNA-seq datasets covering AD patients and non-pathology individuals were obtained from the Gene Expression Omnibus (https://www.ncbi.nlm.nih.gov/geo) as listed: GSE129308^60^, GSE138852^61^, GSE147528^62^, GSE157827^29^, GSE160936^63^, GSE167494^64^, GSE174367^65^. A total of 61 AD samples and 59 normal control samples were collected across four brain regions for analysis, including the prefrontal cortex, entorhinal cortex, superior frontal gyrus, and somatosensory cortex. Count matrices were subjected to Scanpy^66^ (version 1.10.4) for subsequent analyses. To ensure the dataset quality, the calculate_qc_metrics function was applied to check the distribution of the counts of unique molecular identifiers (UMIs) in cells, mitochondrial percentage, and the number of detected genes in cells. The AD6 sample from the dataset GSE157827 was discarded due to the high percentage of counts in mitochondrial genes.

Potential doublets in each sample of each dataset were detected using scDblFinder^67^ (version 1.16.0) with default parameter settings. Samples within the same dataset were then combined into a single matrix for cell quality control.

Low-quality cells were excluded if their UMI counts were greater than 20,000 or less than 200, and the of detected gene number was not between 200 and 8,000. Additionally, cells containing more than 10% mitochondrial DNA were also discarded. Genes present in more than three cells were retained as expressed genes. Then, potential doublets identified by scDblFinder were discarded from the dataset. All processed datasets were combined into a single matrix, which was subject to normalization using the normalize_total function by setting a target sum of 10,000, followed by a logarithmic transformation with the log1p function.

To minimize the batch effect correction, three thousand highly variable genes (HVGs) were selected using the highly_variable_genes function with seurat_v3 flavor, specifying ‘sample’ as the batch_key. scVI^68^ (version 1.3.0) was used to correct the batch effect using the dataset as the batch key and the sample as a categorical covariate. The model was set up with a maximum of 400 epochs, early stopping patience of 10, and a batch size of 2048. The latent embeddings were extracted to the matrix of all features to calculate the nearest neighbor distances for clustering and dimensionality reduction.

With the resolution of 0.10, eight clusters were identified by the Leiden algorithm in Scanpy. The marker genes of following eight cell types documented in ssREAD^69^ database were extracted for annotation: astrocytes, endothelial cells, inhibitory neurons, excitatory neurons, microglia, oligodendrocytes, oligodendrocyte precursor cells (OPCs), and pericytes. All cell types were separated except for endothelial cells and pericytes. Therefore, clusters with high expression of marker genes for these two cell types were labeled as “Endo.Pericytes”. The annotation accuracy was evaluated by comparison with the identity labelled in original datasets using a confusion matrix.

### Differential Expression Analysis

Differentially expressed genes between AD and normal groups were detected using the MAST package (version 1.32.0) ^70^in R on each cell type separately to account for dropout (zero expression) modelling. Genes expressed in at least 5% of cells in any given cell type were retained and subjected to normalization. The cellular detection rate (CDR), calculated as the fraction of detected genes per cell, was also used as a covariate in the model formulation to account for technical variation besides tissue region, sex, and postmortem interval (PMI). Benjamini-Hochberg (BH) approach was applied to control the False Discovery Rate of multiple testing. Significant genes were selected using an adjusted p-value < 0.05, and |logFoldChange| > 0.1 were considered. RG4BPs and their target genes within Differentially expressed genes (DEGs) were then curated.

### Functional Enrichment Analysis

Functional enrichment analysis DEGs and CIRBP targeting DEGs was conducted using the Metascape online tool^7^1 (www.metascape.org). All the expressed genes detected in the single-cell dataset were set as the background. The analysis focused on Gene Ontology Biological Processes, KEGG Pathway, and Reactome Gene Sets. The following threshold was set to select significantly enriched terms: p-value < 0.05, a minimum count of 3, and an enrichment factor > 1.5. To reduce redundancy, significant terms were grouped into clusters based on their similarities using a similarity threshold of 0.3.

### Sub-clustering and transcriptional state identification

#### Astrocyte

One astrocyte snRNA-seq dataset was used to detect distinct transcriptional states ^12^. This dataset sampled astrocytes from five distinct brain regions—the entorhinal cortex (EC), inferior temporal gyrus (ITG), dorsolateral prefrontal cortex (PFC), secondary visual cortex (V2), and primary visual cortex (V1)—collected from 32 donors spanning the spectrum from normal ageing to severe AD. QC was performed according to the original study. Protein-coding genes with more than UMI count 100 and were expressed in at least 30% of samples within each brain region were kept. Cells expressing fewer than 2,000 genes or containing more than 25,000 total UMI counts were excluded. Data after QC were merged into a single matrix consisting of 370,109 cells and 12,838 genes for downstream analysis.

The UMI count matrix was normalized to 10,000 counts per cell and log-transformed (log1p). The top 2,000 HVGs were identified using the “seurat_v3” method in Scanpy. Principal Component Analysis (PCA) was performed based on HVGs. Harmony was applied to the top 50 principal components (PCs) to mitigate batch effects, in which the “SampleName” identifier was specified as a covariate. A neighborhood graph was computed based on the Harmony-integrated representation to conduct dimensionality reduction and clustering. The Louvain algorithm was employed using a resolution of 0.7, yielding 14 initial clusters, which were visualized through Uniform Manifold Approximation and Projection (UMAP). Cluster annotation was checked and confirmed by examining canonical marker genes and calculating the Jaccard index to measure the similarity between the DEGs of each cluster and the cell-type signatures defined in the original study. These 14 clusters were further annotated into six astrocyte transcriptional states: Homeostatic (astH0), Intermediate (astIM), Metabolic (astMet), Reactive 1 (astR1), Reactive 2 (astR2), and astTinf in addition to two putative doublet clusters (astMic and astNeu).

Individuals were grouped into control and AD groups for comparison between healthy individuals and AD patients. To further compare transcriptional dynamics during AD progression, donors were stratified based on the pathology stages defined in the original study ^12^. Individuals in Pathology Stage 1, defined by the absence of neuritic plaques (NPs) and early Braak neurofibrillary tangle (NFT) stages (0–II) representing a “low Alzheimer’s disease neuropathologic change (ADNC) burden”, were defined as the “nonAD” group. Conversely, individuals in Pathology Stages 2–4 were classified as the AD. More specifically, patients in Pathology Stage 2 were defined as “earlyAD”, while those in Pathology Stages 3-4 were categorized as “lateAD”. These stages correspond to “intermediate to high ADNC burden”, exhibiting sparse-to-frequent NPs and advanced Braak NFT stages (ranging from II/III to VI).

DEGs were detected using MAST by including CDR, brain region, and sex as covariates. An absolute log fold change > 0.1 and adjusted p-value < 0.05 was set to obtain differentially expressed gene. Subsequently, functional enrichment analysis was conducted using the clusterProfiler package (version, citation) to elucidate the biological functions and pathways associated with these DEGs, focusing on Gene Ontology.

#### Microglia

One processed microglia snRNA-seq dataset was used to investigate RG4BPs across heterogeneous transcriptional states within microglia ^13^. Data was pre-processed using Seurat, followed by normalization and identification of 2,000 HGVs for PCA. Harmony was applied to the top 30 PCs to correct technical variations by setting “batch” as a covariate. After the construction of K-nearest neighbor (KNN) graph, clusters were identified using Louvain algorithm with a resolution of 0.6. A secondary QC was further applied to exclude brain-associated macrophages (BAMs) (cluster 10), T cells (cluster 13), and other non-microglial cluster (cluster 16) by examining canonical markers. For the remaining microglial clusters, *de novo* marker genes were identified using the FindAllMarkers function (Wilcoxon rank-sum test). Cluster identify was annotated by integrating module scores derived from the original marker lists (calculated via AddModuleScore) and Gene Ontology (GO) enrichment analysis derived from detected *de novo* markers. Clusters were annotated to eight distinct microglia identities, including Homeostatic (cluster 0, 2, and 4), Lipid Processing (cluster 3), Ribosome Biogenesis (cluster 5), Stress Related (cluster 7 and 11), Glycolytic (cluster 8), Antiviral (cluster 9), Inflammatory 1, 2, and 3 (cluster 1, 6, and 15), Cycling (cluster 14), and one Unknown cluster (cluster 11).

Individuals were stratified into control, early AD and late AD groups based on their original diagnosis. The control group comprised individuals diagnosed as “nonAD”, while those classified as “earlyAD” or “lateAD” were combined into the AD group. DEGs were detected using MAST by including following factors as covariates: CDR, batch, brain region, sex, mitochondrial percentage, ribosomal percentage, PMI, and age. DEGs were detected using the following thresholds: adjusted p-value < 0.05 and an absolute log fold change > 0.1. Corresponding gene Ontology (GO) enrichment on DEGs was conducted using clusterProfilers.

### Construction of gene modules

The hdWGCNA^72^ (version 0.4.8) was used to identify RG4BP-associated co-expression gene modules separately for astrocytes and microglia with a customized gene list. This gene list was composed of three sources: the top 2,000 HVGs, expressed RG4BP genes, and the differentially expressed target genes of RG4BP, respectively. Metacells were constructed to aggregate similar cells using the ConstructMetacells function with the KNN algorithm (k = 50 neighbors per metacell) based on the Harmony-integrated PCA reduction. Matrices of metacells were then normalized using the function NormalizeMetacells. A signed network was constructed, followed by selecting a soft-thresholding power to maximize the satisfaction of scale-free topology. Gene modules were identified using a hierarchical clustering approach, in which module expression scores were calculated for single cells using the UCell algorithm (via ModuleExprScore). Module Eigengenes (MEs) that represent the first principal component of gene expression within each module were obtained to interpret potential biological features. Genes with high connectivity within a module were defined as hub genes. Module membership (kME) for each gene, which is defined as the correlation between the gene’s expression and the module eigengene, was caculated. Functional enrichment analysis of the genes within significantly associated modules was performed using the RunEnrichr function to identify associated biological processes (GO-BP) using GO 2025.

To prioritize biologically robust gene interactions, an integrated network by combining transcriptomic co-expression with physical protein-level evidence from the STRING^7^3 database (version 12.0) was constructed. Specifically, an integrated network was generated by masking the hdWGCNA Topological Overlap Matrix (TOM) with binarized protein-protein interaction (PPI) data (confidence score > 200). This step retained co-expression weights only for gene pairs with direct PPI at both transcriptome and proteome-level. Within these integrated networks, dual-evidence hub genes were identified by selecting candidates that ranked in the top 25 for both module membership (kME) and connectivity (degree) within the PPI-validated network. The Cytoscape^74^ was used for the network visualization.

## Data availability

The datasets used in this study are accessible in the GEO or https://compbio.mit.edu/microglia_states ^13^: ref.^60^ GSE129308; ref.^61^ GSE138852; ref.^62^ GSE147528; ref.^29^ GSE157827; ref.^63^ GSE160936; ref.^64^ GSE167494; ref.^65^ GSE174367; ref.^12^ GSE268599; ref.^58^ GSE225516; ref.^57^ GSE164523.

## Acknowledgements

This work was supported by a start-up grant for new faculty (Grant No. 7200735), APRT (Grant No. 9610580), CityU internal grant (Grant No. 7006044 and 9609330) to Jilin Zhang by City University of Hong Kong; National Natural Science Foundation of China (Grant No. 32471343 and 32222089), Research Grants Council of the Hong Kong Special Administrative Region (Grant No. CityU 11101525, Grant No. RFS2425-1S02, Grant No. CityU 11100123, and Grant No. CityU 11100222), Croucher Foundation (Grant No. 9509003), State Key Laboratory of Marine Environmental Health (Grant No. SCRF0070), City University of Hong Kong (Grant No. 9680376, 7030001, and 9678302) to Chun Kit Kwok.

## Contributions

**Xingyu Zhou:** Formal analysis, Investigation, Methodology, Software, Visualization, and Writing – original draft. **Yuanyuan Zhang:** Investigation, Formal analysis, and Software. **Lei Sun:** Validation and Resources, Writing – review & editing. **Chun Kit Kwok:** Funding acquisition, Resources, Validation, Supervision, and Writing – review & editing. **Jilin Zhang:** Conceptualization, Funding acquisition, Methodology, Project administration, Resources, Supervision, Writing – original draft, and Writing – review & editing

